## Supplementary Files for "Human DCP1 is crucial for mRNA decapping and possesses paralog-specific gene regulating functions"

### Fig.S1

A

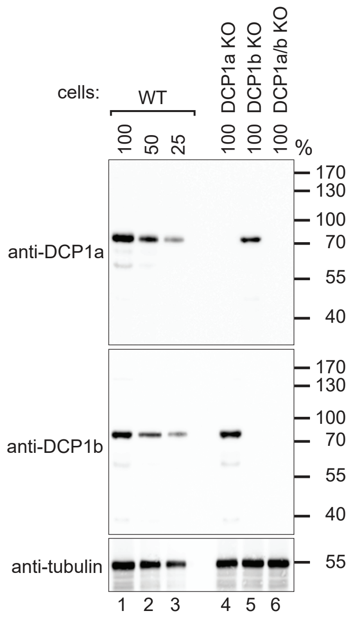


B

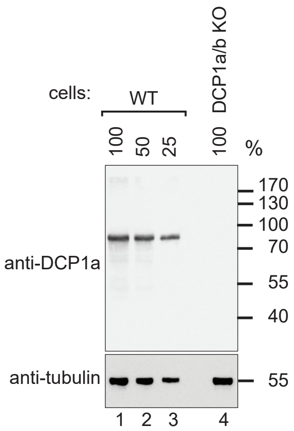


C

**DCP1a/b KO clone:**

***DCP1a:***

```
           WT   AGGTTCCCCCCAGCTTACCTCGAAACAGCACCATGATG
sgRNA sequence           CCAGCTTACCTCGAAACAGC
     allele #1   AGGTTCCCCCCA--TTACCTCGAAACAGCACCATGATG (-2)
     allele #2   AGGTTCCCCCAAG--TACCTCGAAACAGCACCATGATG (-2)
```

***DCP1b:***

```
           WT   TGTACACCTTCGGCCATCGGGCCAACGAGTGGGTGCGT
sgRNA sequence           TCGGCCATCGGGCCAACGAG
     allele #1   TGTACACCTTCGGC-----------CGAGTGGGTGCGT (-11)
     allele #2   TGTACACCTTCGGCCA-----------------TGCGT (-17)
```

#### Fig.S2

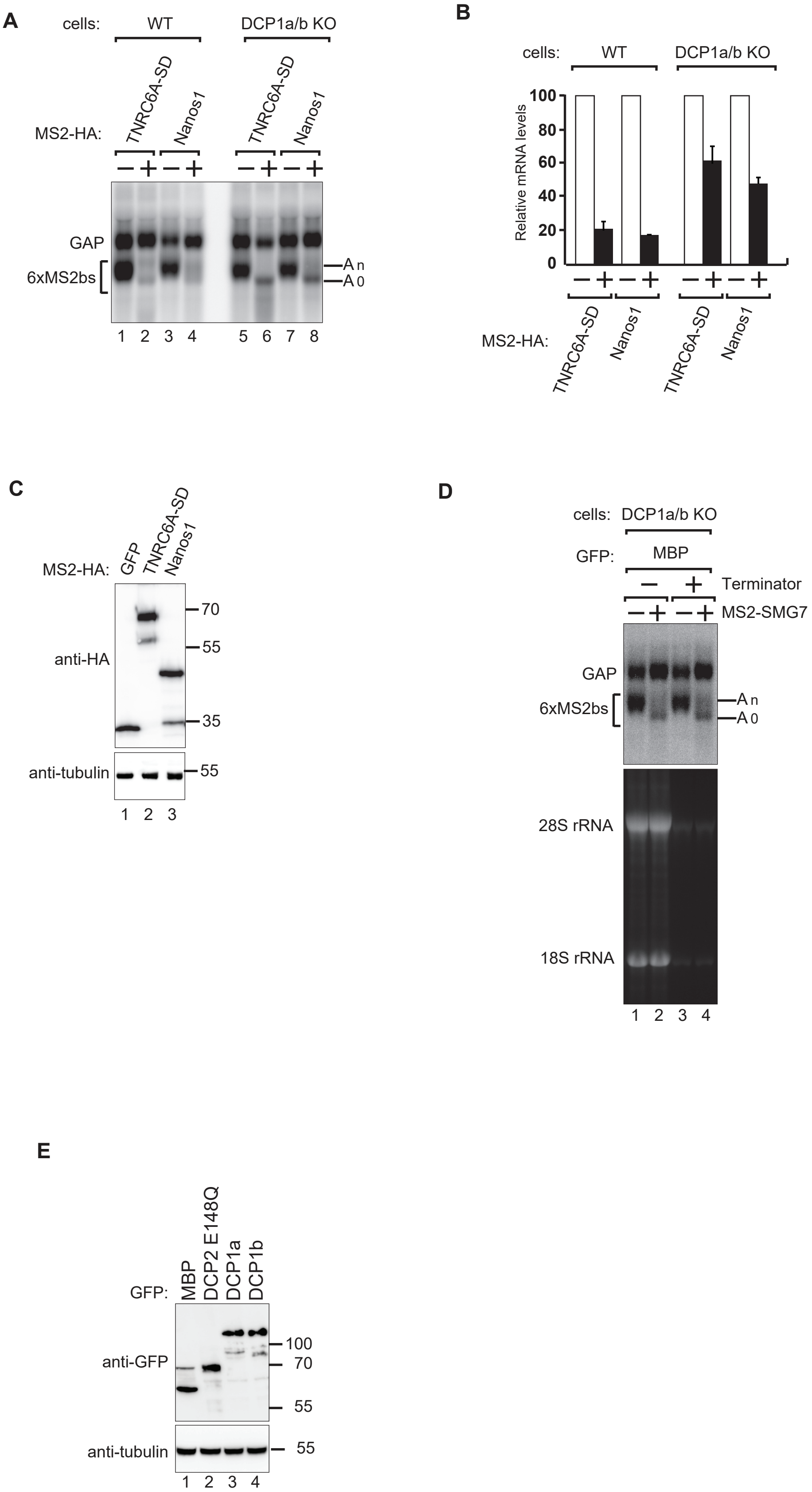
A
cells:
WT
DCP1a/b KO
MS2-HA:
TNRC6A-SD
Nanos1
GAP
6xMS2bs
An
A0
1 2 3 4 5 6 7 8
B
cells:
WT
DCP1a/b KO
Relative mRNA levels
100
80
60
40
20
0
MS2-HA:
TNRC6A-SD
Nanos1
C
MS2-HA:
GFP
TNRC6A-SD
Nanos1
anti-HA
70
55
35
anti-tubulin
55
1 2 3
D
cells: DCP1a/b KO
GFP: MBP
Terminator
MS2-SMG7
GAP
6xMS2bs
An
A0
28S rRNA
18S rRNA
1 2 3 4
E
GFP:
MBP
DCP2 E148Q
DCP1a
DCP1b
anti-GFP
100
70
55
anti-tubulin
55
1 2 3 4

### Fig.S3

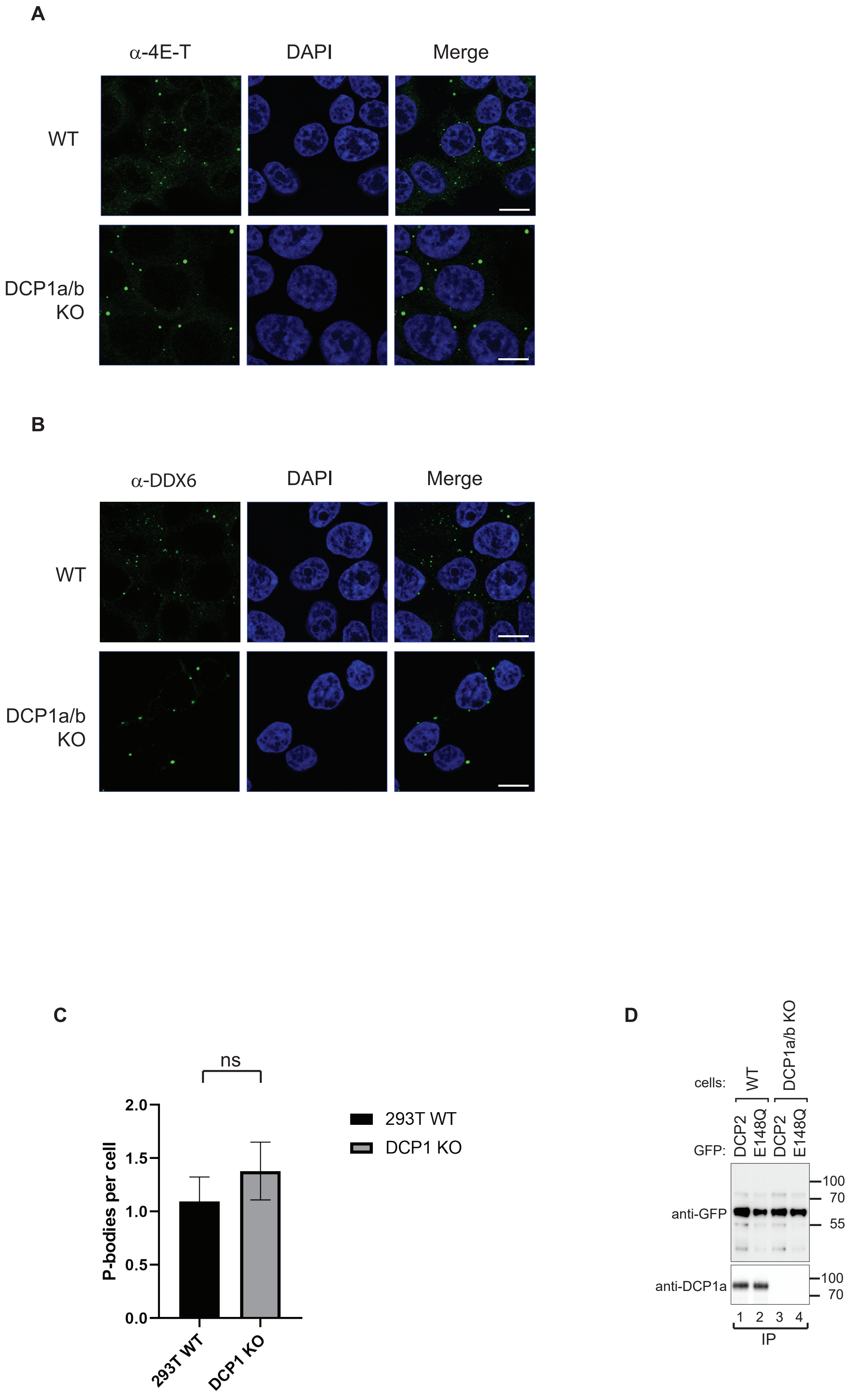
A
α-4E-T
DAPI
Merge
WT
DCP1a/b
KO
B
α-DDX6
DAPI
Merge
WT
DCP1a/b
KO
C
ns
2.0
1.5
1.0
0.5
0.0
P-bodies per cell
293T WT
DCP1 KO
293T WT
DCP1 KO
D
cells:
WT
DCP1a/b KO
GFP:
DCP2
E148Q
DCP2
E148Q
anti-GFP
anti-DCP1a
100
70
55
100
70
1 2 3 4
IP

### Fig.S4

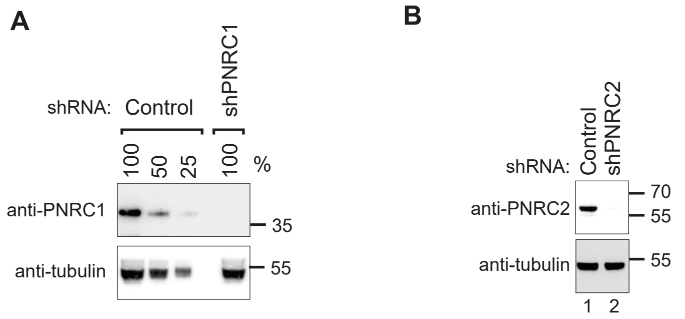
A
shRNA:
Control
shPNRC1
100
50
25
100
%
anti-PNRC1
35
anti-tubulin
55
B
shRNA:
Control
shPNRC2
anti-PNRC2
70
55
anti-tubulin
55
1
2

### Fig.S5

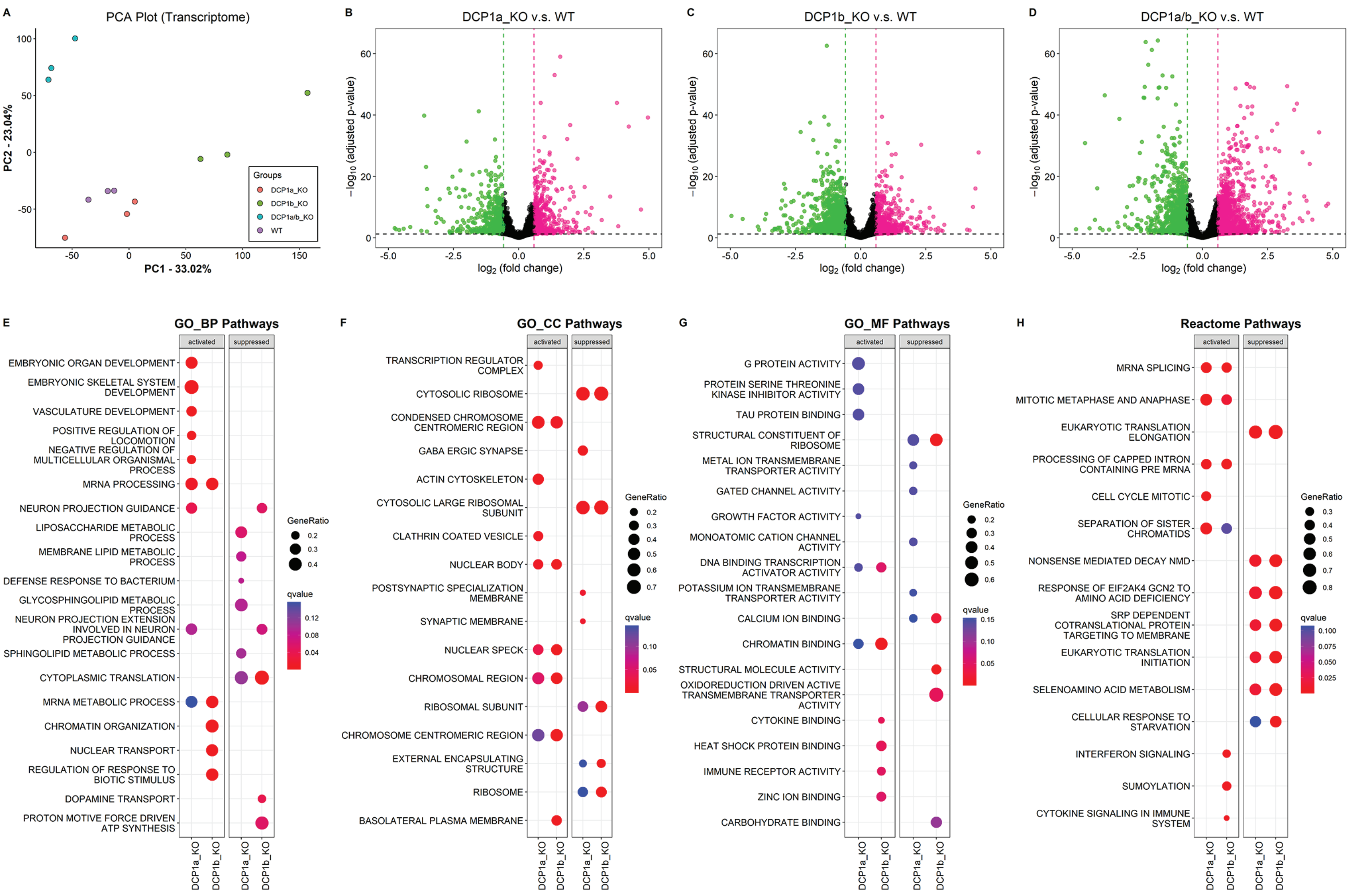
A
PCA Plot (Transcriptome)
PC1 - 33.02%
PC2 - 23.04%
Groups
DCP1a_KO
DCP1b_KO
DCP1a/b_KO
WT
B
DCP1a_KO v.s. WT
C
DCP1b_KO v.s. WT
D
DCP1a/b_KO v.s. WT
$-\log_{10}$ (adjusted p-value)
$\log_2$ (fold change)
E
GO_BP Pathways
activated
suppressed
EMBRYONIC ORGAN DEVELOPMENT
EMBRYONIC SKELETAL SYSTEM DEVELOPMENT
VASCULATURE DEVELOPMENT
POSITIVE REGULATION OF LOCOMOTION
NEGATIVE REGULATION OF MULTICELLULAR ORGANISMAL PROCESS
MRNA PROCESSING
NEURON PROJECTION GUIDANCE
LIPOSACCHARIDE METABOLIC PROCESS
MEMBRANE LIPID METABOLIC PROCESS
DEFENSE RESPONSE TO BACTERIUM
GLYCOSPHINGOLIPID METABOLIC PROCESS
NEURON PROJECTION EXTENSION INVOLVED IN NEURON PROJECTION GUIDANCE
SPHINGOLIPID METABOLIC PROCESS
CYTOPLASMIC TRANSLATION
MRNA METABOLIC PROCESS
CHROMATIN ORGANIZATION
NUCLEAR TRANSPORT
REGULATION OF RESPONSE TO BIOTIC STIMULUS
DOPAMINE TRANSPORT
PROTON MOTIVE FORCE DRIVEN ATP SYNTHESIS
GeneRatio
qvalue
DCP1a_KO
DCP1b_KO
F
GO_CC Pathways
TRANSCRIPTION REGULATOR COMPLEX
CYTOSOLIC RIBOSOME
CONDENSED CHROMOSOME CENTROMERIC REGION
GABA ERGIC SYNAPSE
ACTIN CYTOSKELETON
CYTOSOLIC LARGE RIBOSOMAL SUBUNIT
CLATHRIN COATED VESICLE
NUCLEAR BODY
POSTSYNAPTIC SPECIALIZATION MEMBRANE
SYNAPTIC MEMBRANE
NUCLEAR SPECK
CHROMOSOMAL REGION
RIBOSOMAL SUBUNIT
CHROMOSOME CENTROMERIC REGION
EXTERNAL ENCAPSULATING STRUCTURE
RIBOSOME
BASOLATERAL PLASMA MEMBRANE
G
GO_MF Pathways
G PROTEIN ACTIVITY
PROTEIN SERINE THREONINE KINASE INHIBITOR ACTIVITY
TAU PROTEIN BINDING
STRUCTURAL CONSTITUENT OF RIBOSOME
METAL ION TRANSMEMBRANE TRANSPORTER ACTIVITY
GATED CHANNEL ACTIVITY
GROWTH FACTOR ACTIVITY
MONOATOMIC CATION CHANNEL ACTIVITY
DNA BINDING TRANSCRIPTION ACTIVATOR ACTIVITY
POTASSIUM ION TRANSMEMBRANE TRANSPORTER ACTIVITY
CALCIUM ION BINDING
CHROMATIN BINDING
STRUCTURAL MOLECULE ACTIVITY
OXIDOREDUCTION DRIVEN ACTIVE TRANSMEMBRANE TRANSPORTER ACTIVITY
CYTOKINE BINDING
HEAT SHOCK PROTEIN BINDING
IMMUNE RECEPTOR ACTIVITY
ZINC ION BINDING
CARBOHYDRATE BINDING
H
Reactome Pathways
MRNA SPLICING
MITOTIC METAPHASE AND ANAPHASE
EUKARYOTIC TRANSLATION ELONGATION
PROCESSING OF CAPPED INTRON CONTAINING PRE MRNA
CELL CYCLE MITOTIC
SEPARATION OF SISTER CHROMATIDS
NONSENSE MEDIATED DECAY NMD
RESPONSE OF EIF2AK4 GCN2 TO AMINO ACID DEFICIENCY
SRP DEPENDENT COTRANSLATIONAL PROTEIN TARGETING TO MEMBRANE
EUKARYOTIC TRANSLATION INITIATION
SELENOAMINO ACID METABOLISM
CELLULAR RESPONSE TO STARVATION
INTERFERON SIGNALING
SUMOYLATION
CYTOKINE SIGNALING IN IMMUNE SYSTEM

### Fig.S6

A

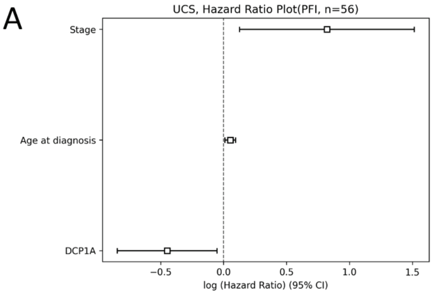


| | HR | Confidence Interval | P-val | PH-assumption |
|---|---|---|---|---|
| Age at diagnosis | 1.05 | [1.01, 1.10] | 1.65E-02 | 0.09 |
| Stage | 2.27 | [1.13, 4.55] | 2.06E-02 | 0.74 |
| DCP1A | 0.64 | [0.43, 0.95] | 2.62E-02 | 0.12 |

*HR : Hazard ratio, TR : time ratio

B

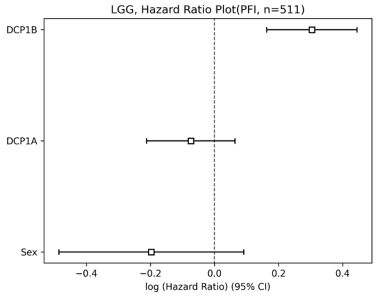


| | HR | Confidence Interval | P-val | PH-assumption |
|---|---|---|---|---|
| DCP1B | 1.36 | [1.18, 1.56] | 2.46E-05 | 0.81 |
| Sex | 0.82 | [0.62, 1.10] | 1.80E-01 | 0.32 |
| DCP1A | 0.93 | [0.81, 1.07] | 2.92E-01 | 0.87 |

*HR : Hazard ratio, TR : time ratio

C

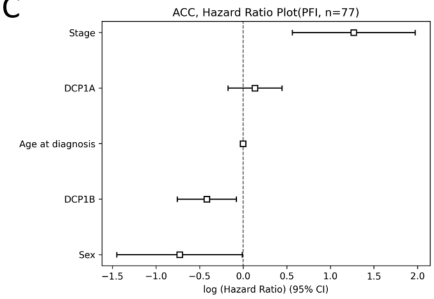


| | HR | Confidence Interval | P-val | PH-assumption |
|---|---|---|---|---|
| Stage | 3.55 | [1.76, 7.19] | 4.21E-04 | 0.94 |
| DCP1B | 0.66 | [0.47, 0.93] | 1.60E-02 | 0.11 |
| Sex | 0.48 | [0.23, 0.99] | 4.69E-02 | 0.08 |
| DCP1A | 1.14 | [0.84, 1.56] | 3.94E-01 | 0.46 |
| Age at diagnosis | 1.00 | [0.98, 1.02] | 8.22E-01 | 0.65 |

*HR : Hazard ratio, TR : time ratio

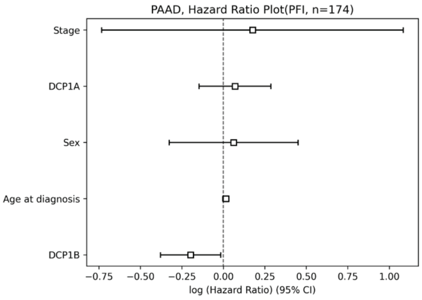


| | HR | Confidence Interval | P-val | PH-assumption |
|---|---|---|---|---|
| DCP1B | 0.82 | [0.68, 0.98] | 3.20E-02 | 0.14 |
| Age at diagnosis | 1.01 | [0.99, 1.03] | 1.52E-01 | 0.80 |
| DCP1A | 1.07 | [0.86, 1.33] | 5.31E-01 | 0.08 |
| Stage | 1.19 | [0.48, 2.96] | 7.06E-01 | 0.41 |
| Sex | 1.06 | [0.72, 1.57] | 7.56E-01 | 0.43 |

*HR : Hazard ratio, TR : time ratio

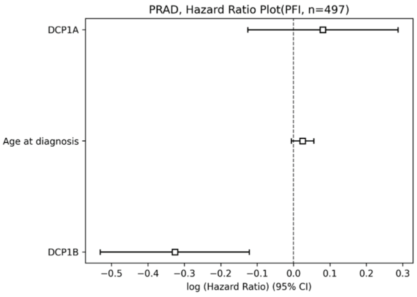


| | HR | Confidence Interval | P-val | PH-assumption |
|---|---|---|---|---|
| DCP1B | 0.72 | [0.59, 0.89] | 1.77E-03 | 0.68 |
| Age at diagnosis | 1.02 | [0.99, 1.06] | 1.20E-01 | 0.35 |
| DCP1A | 1.08 | [0.88, 1.33] | 4.45E-01 | 0.16 |

*HR : Hazard ratio, TR : time ratio

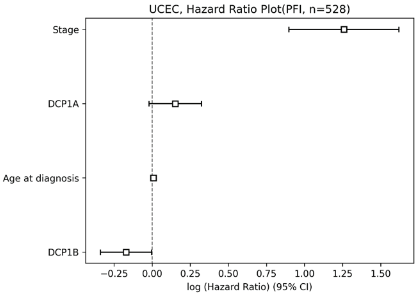


| | HR | Confidence Interval | P-val | PH-assumption |
|---|---|---|---|---|
| Stage | 3.52 | [2.45, 5.05] | 8.34E-12 | 0.81 |
| DCP1B | 0.84 | [0.71, 1.00] | 4.54E-02 | 0.86 |
| DCP1A | 1.16 | [0.98, 1.38] | 8.48E-02 | 0.47 |
| Age at diagnosis | 1.01 | [0.99, 1.03] | 3.29E-01 | 0.09 |

*HR : Hazard ratio, TR : time ratio

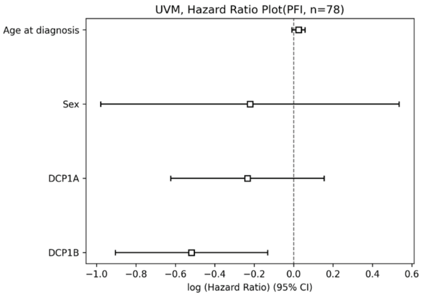


| | HR | Confidence Interval | P-val | PH-assumption |
|---|---|---|---|---|
| DCP1B | 0.60 | [0.40, 0.88] | 8.62E-03 | 0.56 |
| Age at diagnosis | 1.02 | [0.99, 1.06] | 1.48E-01 | 0.30 |
| DCP1A | 0.79 | [0.54, 1.17] | 2.38E-01 | 0.68 |
| Sex | 0.80 | [0.38, 1.71] | 5.66E-01 | 0.94 |

*HR : Hazard ratio, TR : time ratio

### Fig.S7

A

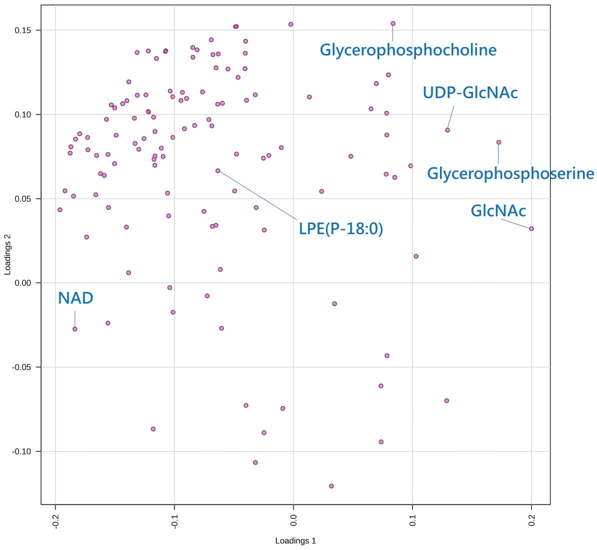

Glycerophosphocholine
UDP-GlcNAc
Glycerophosphoserine
GlcNAc
LPE(P-18:0)
NAD
Loadings 1
Loadings 2


B

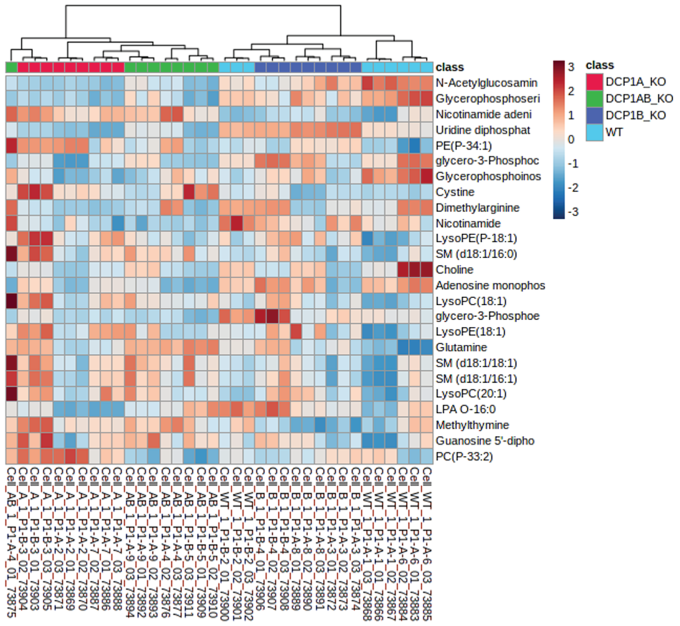

class
N-Acetylglucosamin
Glycerophosphoseri
Nicotinamide adeni
Uridine diphosphat
PE(P-34:1)
glycero-3-Phosphoc
Glycerophosphoinos
Cystine
Dimethylarginine
Nicotinamide
LysoPE(P-18:1)
SM (d18:1/16:0)
Choline
Adenosine monophos
LysoPC(18:1)
glycero-3-Phosphoe
LysoPE(18:1)
Glutamine
SM (d18:1/18:1)
SM (d18:1/16:1)
LysoPC(20:1)
LPA O-16:0
Methylthymine
Guanosine 5'-dipho
PC(P-33:2)
class
DCP1A_KO
DCP1AB_KO
DCP1B_KO
WT


C

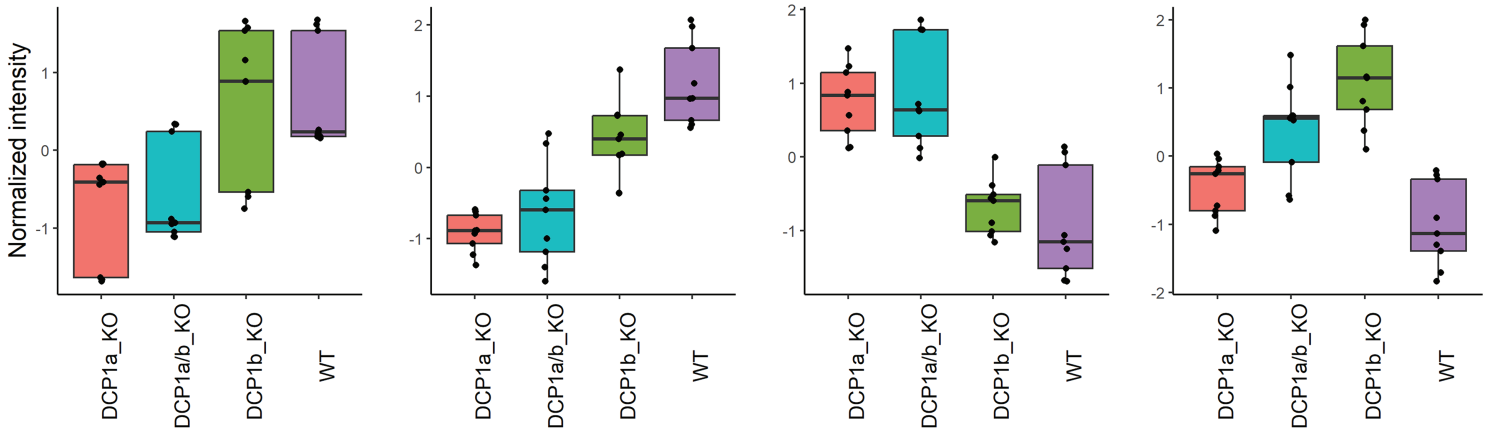

Glycerophosphocholine
Glycerophosphoserine
NAD
LPE(P-18:0)
Normalized intensity
DCP1a_KO
DCP1a/b_KO
DCP1b_KO
WT

### Fig.S8

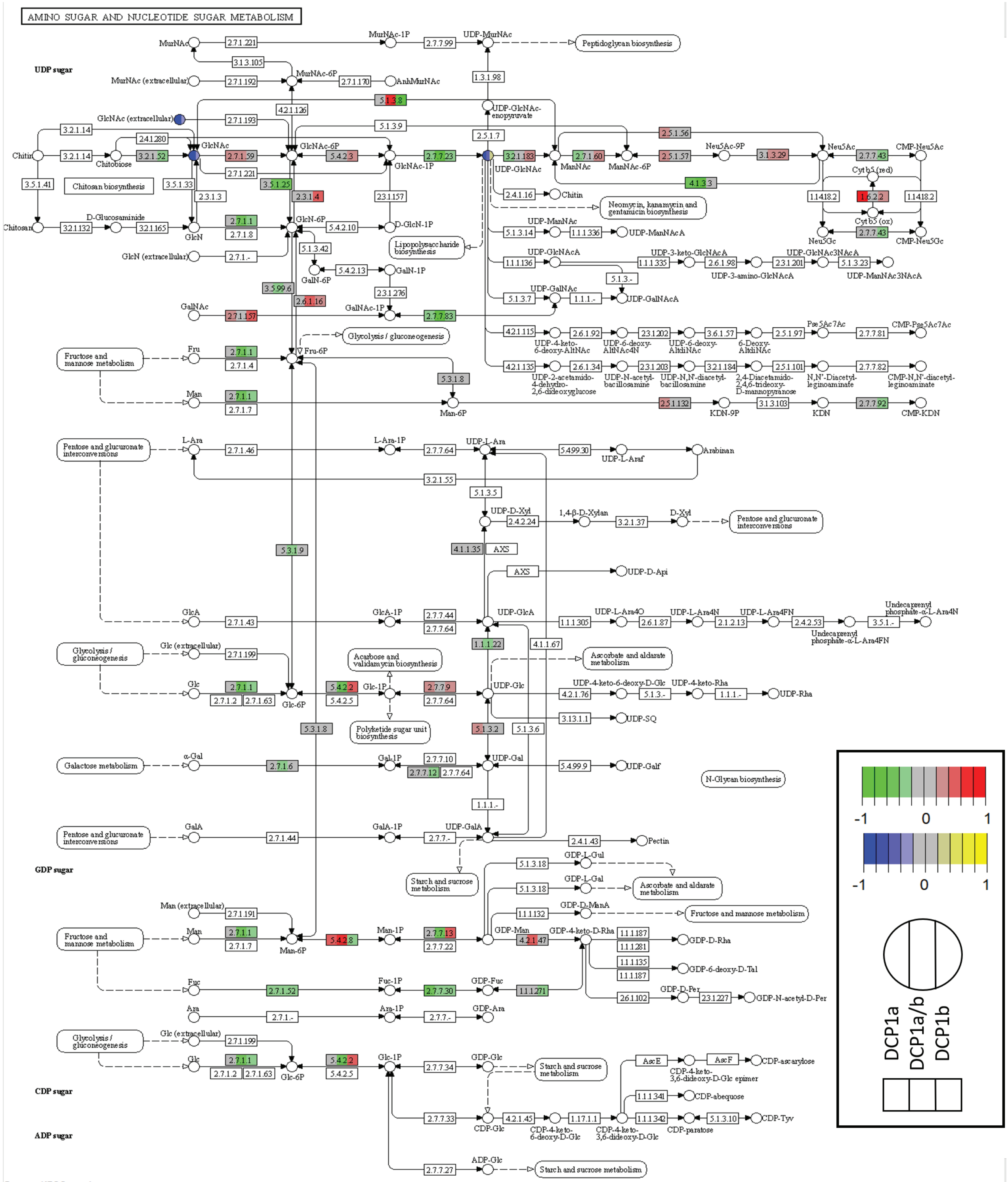


Data on KEGG graph
Rendered by Pathview

**Supplementary Table S1** Constracts used in this study.

| Name | *Hs* DCP2 | Plasmid |
|---|---|---|
| DCP2 | 1–420 | pT7-EGFP-DCP2,<br>pCIneo-V5-SBP-DCP2 |
| E148Q | 1–420 (E148Q) | pT7-EGFP-DCP2 E148Q |

| Name | *Hs* DCP1a | Plasmid |
|---|---|---|
| DCP1a | 1–582 | pT7-EGFP-DCP1a |
| ΔEVH1 | 132-582 | pT7-EGFP-DCP1a 132-682 |
| ΔHLM | Δ155-168 | pT7-EGFP-DCP1a Δ155-168 |
| ΔTD | 1-538 | pT7-EGFP-DCP1a 1-538 |
| EVH1 | 1-132 | pT7-EGFP-DCP1a 1-132 |
| 1-168 | 1-168 | pT7-EGFP-DCP1a 1-168 |
| 1-254 | 1-254 | pT7-EGFP-DCP1a 1-254 |
| 1-538 | 1-538 | pT7-EGFP-DCP1a 1-538 |

| Name | *Hs* DCP1b | Plasmid |
|---|---|---|
| DCP1b | 1-618 | pT7-EGFP-DCP1b |

| Name | *Hs* PNRC1 | Plasmid |
|---|---|---|
| PNRC1 | 1–327 | pEF-DEST51 HA-PNRC1<br>(Addgene #123294) |

| Name | *Hs* PNRC2 | Plasmid |
|---|---|---|
| PNRC2 | 1–139 | pCIneo-V5-SBP-MBP-PNRC2 |

| Name | *Target gene* | Plasmid |
|---|---|---|
| shPNRC1 | *Hs* PNRC1 | pSUPER.puro-PNRC1#4 |
| shPNRC2 | *Hs* PNRC2 | pSUPER.puro-PNRC2-t2 |

**Supplementary Table S2** Antibodies used in this study.

| Antibody | Source | Catalog Number | Dilution | Monoclonal/ Polyclonal |
|---|---|---|---|---|
| Anti-V5 | AbD Serotec | MCA1360GA | 1:5,000 | Mouse Monoclonal |
| Anti-HA-HRP | Roche | 12 013 819 001 | 1:5,000 | Mouse Monoclonal |
| Anti-GFP (for Western blotting) | Roche | 11 814 460 001 | 1:2,000 | Mouse Monoclonal |
| Anti-GFP (for immunoprecipitations) | In house | | | Rabbit Polyclonal |
| Anti-tubulin | Sigma | T6199 | 1:5,000 | Mouse Monoclonal |
| Anti-EDC4 | Santa Cruz Biotechnology | sc-8418 | 1:1,000 | Mouse Monoclonal |
| Anti-DCP2 | Bethyl | A302-597A | 1:1,000 | Rabbit Polyclonal |
| Anti-DCP1a (C-terminal region) | Sigma | D5444 | 1:1,000 | Rabbit Polyclonal (Epitope: a.a 512-528) |
| Anti-DCP1a (N-terminal region) | aviva | ARP39353_T100 | 1:1000 | Rabbit Polyclonal (Epitope: a.a 1-50) |
| Anti-DCP1b | Novus | NBP1-82018 | 1:1,000 | Rabbit Polyclonal (Epitope: a.a 132-236) |
| Anti-EDC3 | Abcam | Ab57780 | 1:1,000 | Mouse Monoclonal |
| Anti-DDX6 | Bethyl | A300-461A | 1:1,000 | Rabbit Polyclonal |
| Anti-PatL1 | Bethyl | A303-482A-M | 1:1,000 | Rabbit Polyclonal |
| Anti-4ET | Bethyl | A300-706A | 1:1,000 | Rabbit Polyclonal |
| Anti-mouse-HRP | GE Healthcare | NA931V | 1:10,000 | Polyclonal |
| Anti-rabbit-HRP | GE Healthcare | NA934V | 1:10,000 | Polyclonal |
